# Repeated administration of pharmaceutical-grade medium chain triglycerides, a common pharmacologic excipient, confers dose-dependent toxicity by the intraperitoneal but not oral route in mice*

**DOI:** 10.1101/2024.11.24.625115

**Authors:** Mitzi Adler-Wachter, Julia Y. Tsai, Brendan N. Schweitzer, Ashley McDonough, Jessica M. Snyder, Melissa Barker-Haliski, Jonathan R. Weinstein

## Abstract

Pharmaceutical-grade medium chain triglycerides (MCTs) are common excipients for in vivo pharmacological studies in laboratory animals, and as an experimental therapeutic in certain metabolic and neurological disorders. In this study, we examined the tolerability of repeated administration of a pharmaceutical-grade formulation of three MCTs—caprylic, capric, and lauric acid - in mice via the oral (PO) and intraperitoneal (IP) routes. We administered either 8 or 4 µL of 100% MCTs or saline/gram of body weight twice daily for seven days. During administration and for seven days after, we monitored weight change and clinical presentation. On day 14, or upon meeting euthanasia criteria, animals were sacrificed for gross necropsy, histology, and complete blood count. We observed significant weight loss, clinical decline and 100% mortality in animals receiving 8 µL/g of MCTs via the IP route of administration. Gross necropsy revealed serosanguinous fluid in the thoracic cavity, dark red mottled lungs, and adhesions in the abdominal cavity. Histology confirmed inflammation of the lungs, mediastinum, and peritoneum. Mild gross lesions and initial weight loss (through day 3) were also present in mice receiving 4 µL/g of MCTs IP. However, these animals regained weight by day seven and exhibited no clinical decline or mortality. None of these adverse effects were seen in animals receiving either 8 µL/g of MCTs PO or 8 µL/g of saline IP. These findings suggest repeated IP administration of MCTs may cause dose-dependent toxicity, and mortality at high doses, but confers no adverse effects when administered via the PO route.

**SIGNIFICANCE STATEMENT:** Medium chain triglycerides (MCTs) are commonly used as an excipient in pharmacological studies involving laboratory animals. Our work provides much needed safety information regarding adverse effects of repeated MCTs administration via the intraperitoneal, but not the oral, route in mice.

## INTRODUCTION

Medium chain triglyceride formulations (MCTs) are common formulation vehicles for hydrophobic investigational compounds in preclinical studies. For example, Miglyol 812, a derivative of coconut and palm oil, is a pharmaceutical- or commercial-grade excipient mixture of several MCTs (Singh Negi, 2019), including 50-65% caprylic acid, which is included in the United States Food and Drug Administration’s (FDA) list of generally recognized safe compounds (Select Committee on GRAS Substances, 2020). Another 30-45% of Miglyol is made up of capric acid, and <2% is lauric acid (Buss, 2018). Combining MCTs in these pharmaceutical-grade formulations improves solubility potential (Singh Negi, 2019). Based on studies in rats and monkeys, pharmaceutical-grade excipients, such as Miglyol, are recommended by the U.S. FDA as an injectable excipient for the Type 2 diabetes therapeutic BYDUREON BCISE in humans (Center for Drug Evaluation and Research, application number 209210Orig1s000). MCTs are generally accepted as safe in animal models when administered via the oral (PO) route (Buss, 2018; Traul, 1999), though some adverse reactions have been noted. Sellers et al. examined the effects of four weeks of daily PO administration of 10 µL of 100% Miglyol per gram of body weight (BW) in Wistar Han IGS rats. Treated rats presented with decreased BW gain, and soft and/or mucoid stools. Necropsy revealed alveolar histiocytosis and focal interstitial inflammation in the lungs. However, these intestinal and BW effects were reversible during a 4-week recovery period (Sellers, 2008). Healing et al. reported that 28-days of daily PO dosing with 10 µL/g of 33% Miglyol produced no adverse reactions in rats, while 30% hydroxypropyl-*B*-cyclodextrin, 20% oleic acid, and 40% tetraethylene glycol each produced minor adverse effects (Healing, 2016). Comparatively, Buss et al saw no adverse effects during weekly subcutaneous administration of smaller volumes (0.15 µL/g) relative to body weight of Miglyol in cynomolgus monkeys, nor in single-dose PO administration of 10 µL/g Miglyol in rats (Buss, 2018), suggesting dose-, administration frequency- and route-dependent outcome variability. Despite these studies examining effects of MCT formulations in other animal models, there is limited literature evaluating the safety of sub-chronic or repeated administration of these excipients in mice. There is a similar lack of data regarding outcomes and/or median lethal dose (LD_50_) of MCT formulations, like Miglyol, administered via the intraperitoneal (IP) route, which is commonly used in pharmacology studies.

Short course administration of IP MCTs as formulation vehicle have not been previously reported to be associated with unfavorable reactions (Bunaciu, 2011; Robaina Cabrera, 2021). However, our group observed adverse reactions in male C57BL/6J mice following twice daily IP injections of pharmaceutical-grade MCTs for seven days at a dose of 8 uL per gram BW. This was an incidental observation during testing of an experimental pharmacologic agent using pharmaceutical-grade MCTs as a vehicle, in which animals receiving vehicle alone exhibited decline. One to seven days following the end of treatment, we observed clinical decline and significant mortality in animals, regardless of treatment group (MCTs with drug versus MCTs alone). We observed highly fluctuating BW throughout the seven-day treatment period, followed by rapid onset of immobility, hunching, piloerection, increased respiratory effort, and mortality in 50-100% of animals. Necropsy revealed variably severe lung hemorrhage and pleural effusion, adhesion of peritoneal organs, and cystic lesions filled with oily fluid around the administration sites.

These observations, in combination with findings by other groups (Sellers, 2008; Healing, 2016; Buss, 2018), suggested that pharmaceutical-grade MCTs may cause organ toxicity in IP administration paradigms, but not via other routes. Because of the relevance to pharmaceutical sciences research, it is important to further characterize and discern how to mitigate potential adverse effects of repeated IP pharmaceutical-grade MCTs administration. In this study we explored the outcome of high- (8 µL/g BW, ∼7.56 g/kg) versus low-(4 µL/g BW, ∼3.78 g/kg) doses of pharmaceutical-grade MCTs when administered via the IP route twice daily (b.i.d.) for seven days. We compared this to high-dose (8 µL/g) PO administration of pharmaceutical-grade MCTs, and IP administration of saline, again at 8 µL/g BW. In doing so, we evaluated the safety of repeated pharmaceutical-grade MCTs administration via two common administration routes. All data presented here were collected for the purpose of this study, and no animals received any compound other than 100% pharmaceutical-grade MCTs or sterile saline. Group sizes were designated based on the needs of the present toxicology study. Our findings suggest that repeated IP pharmaceutical-grade MCTs administration to mice may confer significant dose-dependent adverse effects on clinical presentation and organ toxicity.

## METHODS

### Animals

16- to 20-week-old male C57BL/6J (Strain #000664; *n* = 40) mice were obtained from Jackson Laboratories (Sacramento, CA). Animals were housed in cages of five, and care followed standard housing protocols for University of Washington as described (Meeker, 2019). Animals were acclimated for at least one week following arrival prior to experimental manipulation. At the end of the study or upon meeting euthanasia criteria, animals were euthanized using asphyxiation via gradual displacement with CO_2_. Euthanasia criteria were >30% body weight loss and/or a clinical score of 8 or higher out of 12 (see <u>Health Assessment</u> methods for scoring details). All procedures were approved by University of Washington Institutional Animal Care and Use Committee, in accordance with the National Research Council’s Guide for the Care and Use of Laboratory Animals (National Academy of Sciences, 2011).

### Administration

Administration of 100% current good manufacturing practice (cGMP) grade Miglyol 812 (hereafter referred to as “pharmaceutical-grade MCTs”, Caprylic/capric Triglyceride; product code = C3465; pH = 5; Spectrum Chemical MFG corporation, New Brunswick, NJ) was based on our group’s protocol for experimental drug administration (Lee, 2024). Accordingly, animals received b.i.d. IP or PO pharmaceutical-grade MCTs or sterile saline (Hospira Inc, Lake Forest, IL) for 7 days. Prior to administration, pharmaceutical-grade MCTs was aliquoted into sterile endotoxin-free vials in a biosafety cabinet. Aliquots were stirred for 2 minutes using sterile, endotoxin-free stir bars and allowed to cool to room temperature before administration. Animal weights and clinical scores were collected at time of administration throughout the seven-day experiment, and for an additional seven days following final treatment.

Animals were randomly distributed into 4 treatment groups (**Figure 1**) of 10 animals each (based on standard power analysis which yields 80% power to detect a 50% difference in experimental outcomes between treatment groups with an alpha of 0.05). Our control group received IP 8 µL of sterile saline per gram BW (8 µL S IP; *n* = 10) to assess effects of twice-daily IP administration. The high dose group received IP administration of 8 µL of pharmaceutical-grade MCTs per gram BW (∼7.56 g/kg; 8 µL M IP group). One animal was excluded due to premature decline. Upon veterinary evaluation, this decline was presumed unrelated to MCT administration (*n* = 9 for this group). The low dose group received IP administration of 4 µL of pharmaceutical-grade MCTs per gram BW (∼3.78 g/kg, “low dose”; 4 µL M IP group; *n* = 10). Finally, the PO group received 8 µL of pharmaceutical-grade MCTs per gram BW via oral gavage (8 µL M PO group). Three 8 µL M PO animals were excluded due to gavage-related injury in early PO dosing (*n* = 7 for this group).

**Figure 1:**
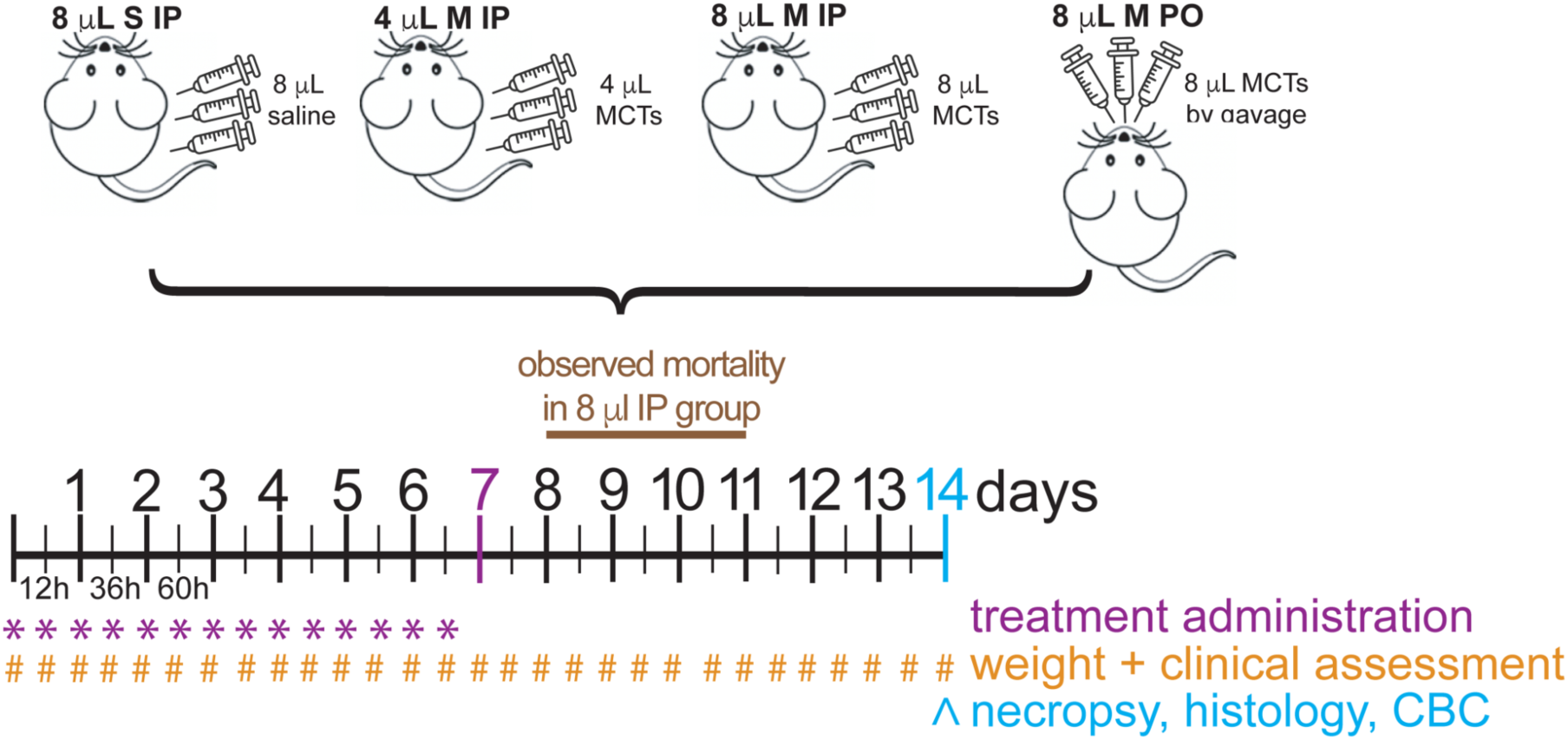
Experimental Design. Animals were split into four groups. The 8 µL S IP group received IP administration of 8 µL sterile saline per gram of BW; the 8 µL M IP group received IP administration of 8 µL pharmaceutical-grade MCTs per gram of BW (7.56 g/kg); the 4 µL M IP group received IP delivery of 4 µL pharmaceutical-grade MCTs per gram of BW (3.78 g/kg); and the 8 µL M PO group received PO boluses of 8 µL pharmaceutical-grade MCTs per gram of BW (7.56 g/kg). All groups received administration of their respective treatments every 12 hours for 7 days. Animals were then monitored, with weight collection and clinical presentation assessment, every 12 hours for an additional 7 days. All remaining animals were euthanized at day 14. Five animals per group in the 8 µL M IP, 4 µL M IP, and 8 µL M PO groups, and two animals in the 8 µL S IP control group were sent for necropsy, histology, and blood collection at time of death or euthanasia.

### Health Assessment

Clinical scores during twice daily health assessments were determined based on criteria in **Table 1**.

**Table 1:** Health assessment criteria.

| Score |  |  |  |  |
| --- | --- | --- | --- | --- |
|  | **0 (normal)** | **1** | **2** | **3 (critical)** |
| Appearance | Normal | Lack of grooming. Mild dehydration.<br>Mild hunched posture. | Moderate piloerection, dehydration** and hunching. Dull eyes. | Significant piloerection and hunching.<br>Sunken eyes.<br>Pale digits. |
| Clinical signs | None | Minor increase in respiratory effort. | Moderately increased respiratory effort. | Rapid respiration. Cold to touch.<br>Abdominal swelling. |
| Behavior | Normal | Mild lethargy but still responsive to stimulation. | Significant lethargy.<br>Decreased activity and response to stimulation. | Immobile.<br>Unresponsive to stimulation.<br>Head bobbing. |
| Body condition* | 3 | 2+ to 3- | 1+ to 2- | 1 |

\*Body Condition Scoring assessment as established in Burkholder et al. (Burkholder, 2012).
Clinical score is a summation of an animal's 0-3 score on each of these four criteria (maximum score = 12).

### Necropsy, histology and blood work

Following euthanasia, five animals each from the 8 µL M IP, 4 µL M IP, and 8 µL M PO groups and two animals from the 8 µL S IP group were necropsied, with tissue collection for histology. Numbers of animals and tissues included for histology were based on severe necropsy and histology findings in the thoracic cavity, peritoneal cavity, and peri-reproductive fat in previous studies using 8 µL pharmaceutical-grade MCTs/g BW. Blood was collected for complete blood count (CBC) from three animals in the 8 µL M IP group, four in the 4 µL M IP group, five in the 8 µL M PO group, and two in the 8 µL S IP group. Two animals in the 8 µL M IP group, and one in the 4 µL M IP group were excluded from CBC due to insufficient blood collection. Necropsy, histology, and blood work were performed by the University of Washington Department of Comparative Medicine Veterinary Diagnostic Laboratory. Briefly, tissues were fixed in 10% neutral buffered formalin and routinely paraffin embedded. Tissue sections (4-5 microns) of the lungs and mediastinal fat, liver, kidneys, heart, pancreas, spleen, mediastinal fat, and male reproductive organs and bladder were stained with hematoxylin and eosin (H&E) and evaluated by a board-certified veterinary pathologist (J.M.S.). Lesions including lung hemorrhage and lung inflammation were scored on a scale of 0-4, with 0 representing normal tissue; 1 representing minimal disease; 2 representing mild disease; 3 representing moderate disease; and 4 representing severe disease. Lesions including inflammation of the mediastinal fat and peritonitis were scored on a scale of 0-2 with 0 representing normal tissue; 1 representing mild disease; and 2 representing moderate to severe disease. Further details on the scoring system are presented in **Supplemental Table 1**. Histology images were captured from glass slides using NIS-Elements BR 4.20.01 and images of gross and histologic lesions were plated in Adobe Photoshop 2023. Image white balance, contrast and lighting were adjusted using auto-corrections applied to the entire image.

### Statistics

Group mean and standard deviation were calculated for weight change and clinical presentation data at each time point and plotted alongside data points for individual animals as comparison. Repeated measure two-way ANOVAs were used to compare the 4 µL M IP and 8 µL M PO groups to the 8 µL S IP control group. A mixed effect model two-way ANOVA was used to compare the 8 µL M IP group to the 8 µL S IP control group. A repeated measure ANOVA for comparison in this case was not possible due to an *n* of 0 in the 8µL M IP group at later time points (resulting from animal deaths in this group). CBC results were averaged by treatment group for each blood work parameter and analyzed by two-way ANOVA. Survival data for each group was calculated as a percentage of original animals that remained alive at each time point and represented as a Kaplan Meier graph and significance was calculated using the Log-Rank test. Semi-quantitative histological scores were analyzed using a Kruskal Wallis Test for non-parametric data with Dunn’s multiple comparisons performed between the 8 µL/g M IP group and all other groups. All statistical analyses and standard deviation statistics were carried out in GraphPad Prism 10.3.1 (Domatics software) and comparisons where p < 0.05 were considered significant. Graphical representations of weight change, clinical score, and survival were created using Python 3.0.

## RESULTS

### Weight Change

In the first three days of administration, animals in the 8 µL M IP and 4 µL M IP groups exhibited a sharp BW decline (**Figure 2**) to an average of 85.6 ± 1.5% and 85.1 ± 1.9% of starting weight (SW), respectively. Animals in the 4 µL M IP group steadily regained weight through study completion, reaching 94.8 ± 2.4% of SW on average at day 14. Mean BW in the 8 µL M IP group continued to slowly decrease, began rebounding on days 6 (86.9 ± 2.9% of SW) and 8 (92.1 ± 3.0% of SW), before sharply declining again to 85.4 ± 4.5% of SW on day 10, as animals died or were euthanized (*n* on day 11 = 1). Weight change for these two groups was significantly different than that of the 8 µL S IP control group, which remained stable and close to SW throughout the 14-day period. Animals in the 8 µL M PO group showed less weight loss, dipping to 94.5 ± 2.3% of SW on day 4. This was followed by a steady BW increase up to 98.4 ± 3.6% of SW on day 9, at which point BW remained stable through endpoint on day 14. Weight change in this group was not significantly different than that of the 8 µL S IP control group (p > 0.05).

**Figure 2:**
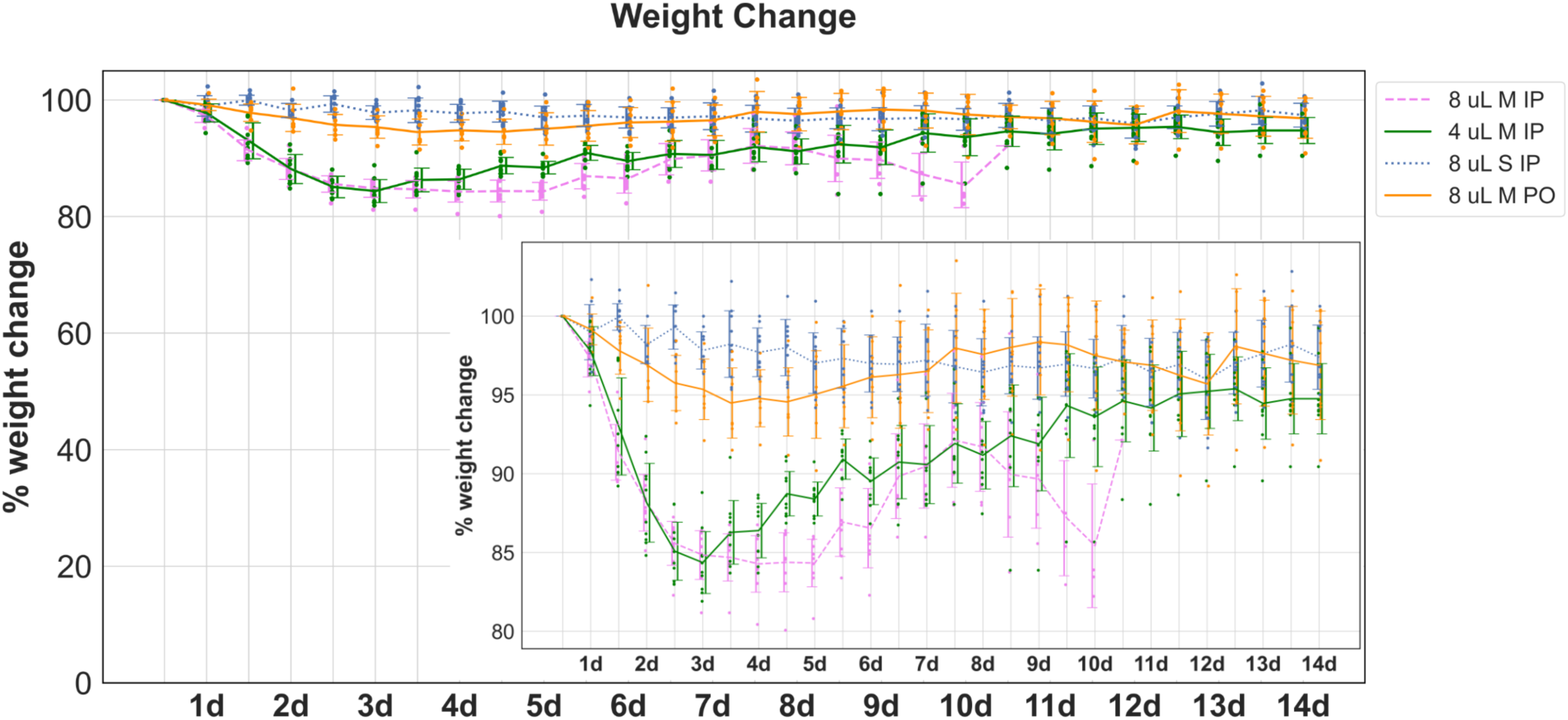
Weight Change. Animals’ body weight was recorded twice daily for 7 days at time of treatment administration and continued for an additional 7 days after the end of the treatment period. Weight change was then calculated as a percentage of starting weight (SW). No animals reached 30% weight loss threshold set as criteria for euthanasia. 8 µL S IP animals weight remained relatively unchanged. 8 µL M PO animals showed moderate, intermittently weight loss (down to 94.5 ± 2.3% of SW) between day 2 and day 5 before returning to around SW and stabilizing by day 8. Both 8 µL M IP and 4 µL M IP groups showed steep weight loss (∼15%) between day 1 and 3. 4 µL M IP animals steadily regained weight, up to ∼94.8 ± 2.4% of SW, meanwhile 8 µL M IP animals more sharply regained weight up to ∼92.1 ± 3.0% of SW between day 6 and 8 at which point weight sharply decreased until day 10. Note: mortalities in the 8 µL M IP group began on day 8 and all animals had died by day 11. Weight change curves for 8 µL M IP and 4 µL M IP groups, but not the 8 µL M PO group, were significantly (p > 0.0001) different than the 8 µL S IP group. 8 µL M IP, day 1 *n* = 9, day 11 *n* = 0; 4 µL M IP, *n* = 10; 8 µL S IP, *n* = 10; 8 µL M PO, *n* = 7.

### Clinical Presentation Score

We also wanted to visually assess clinical presentation during the 14-day monitoring period (**Figure 3**). Scoring was based on criteria in **Table 1**, where scores (0-3) for each of four criteria were totaled to a composite score between 0 (no observed clinical decline) and 12 (severe decline). 8 µL S IP animals presented with no adverse clinical signs, and all animals had a clinical score of 0 throughout the study. Clinical score for the 8 µL M IP group moderately increased (worse presentation) between days 3 and 7, with an average score of up to 1. After the completion of Miglyol administration on day 7, animals in this group began rapidly declining and presented with hunched posture, piloerection, severe lethargy, pale digits, and increased respiratory effort. Because of these symptoms, this group’s clinical score rapidly increased by ∼2 points per day between day 7 and 11, when the last remaining animal was euthanized per exclusion criteria (clinical score > 8). Clinical scores in this group were significantly (p < 0.001) higher than those of the 8 µL S IP.

**Figure 3:**
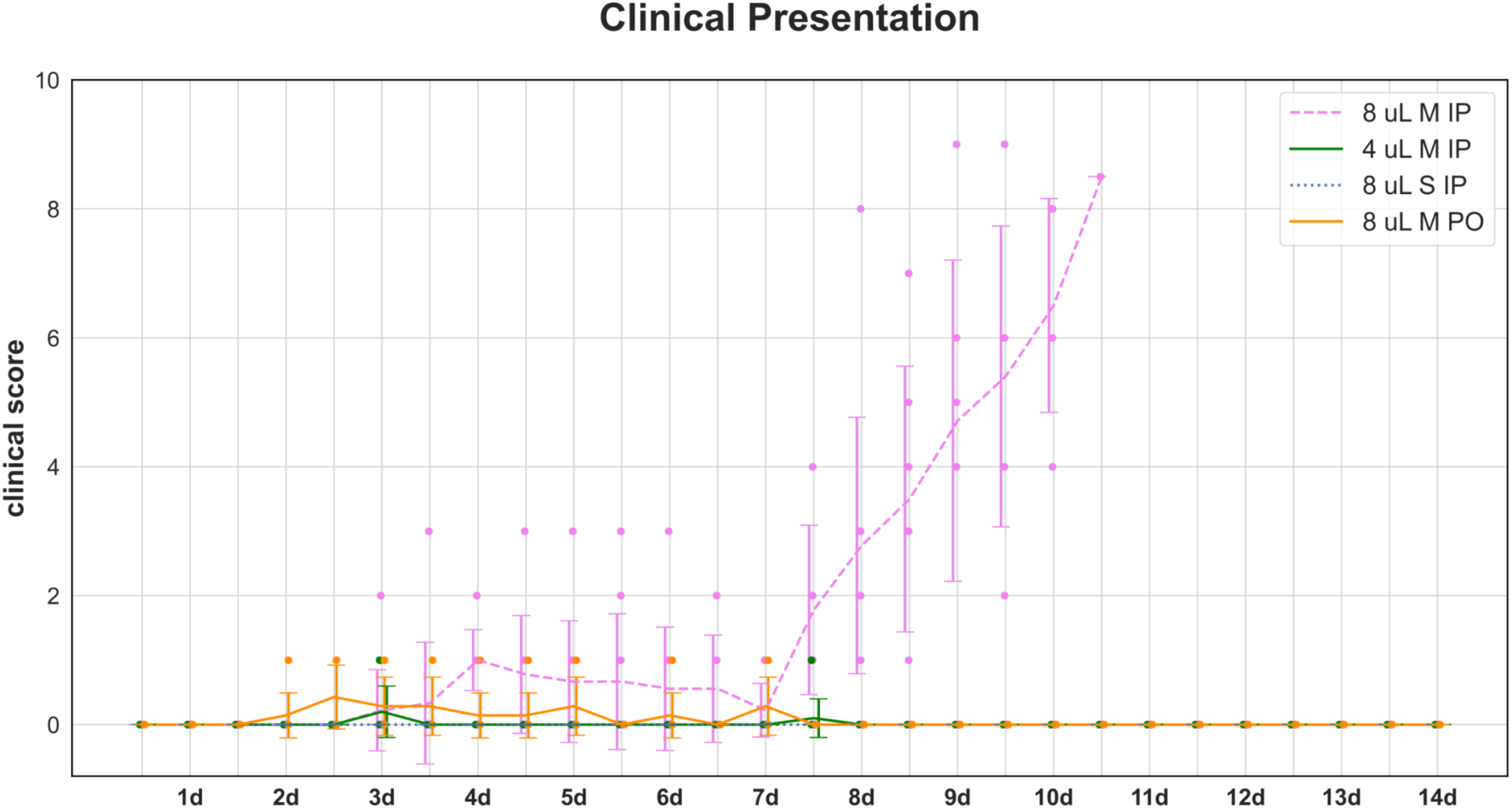
Clinical Presentation. Clinical presentation was assessed based on scoring criteria outlined in **Table 1** twice daily for 7 days at time of treatment administration and continued for an additional 7 days after the end of the treatment period. Appearance, clinical signs of decline, behavior, and body condition each received a score between 0 (normal) and 3 (severe pathology). Total score out of 12 is presented here. 8 µL S IP animals had no change in clinical presentation. Average clinical score for the 8 µL M PO group increased slightly between day 2 and day 7 before returning to baseline on day 8 through 14. Average clinical score for the 4 µL M IP group increased slightly on day 3 and day 8. Average clinical score for animals in the 8 µL M IP group increased to ∼1 between day 3 and day 7, after which point average score sharply increased between day 8 and 10, for a maximum individual score of 9. As compared to the 8 µL S IP group, clinical score for the 8 µL M IP and 8 µL M PO groups were significantly different with p < 0.0001 and p = 0.0005, respectively. Clinical score in the 4 µL M IP group was not significantly different than the 8 µL S IP group. Note: mortalities in the 8 µL M IP group began on day 8 and all animals had died by day 11. 8 µL M IP, day 1 *n* = 9, day 11 *n* = 0; 4 µL M IP, *n* = 10; 8 µL S IP, *n* = 10; 8 µL M PO, *n* = 7.

5/7 animals in the 8 µL M PO demonstrated minor, but significant (p < 0.001), increases in clinical score due to mild to moderate dehydration between day 2 and day 7. Similar mild dehydration was present but not significant in a smaller number of animals (3/10) in the 4 µL M IP group. Both groups remained stable with a score of 0 from day 8 through 14.

### Survival

We also observed significant mortality stratified by treatment period (**Figure 4**). No animals succumbed during the 2X/daily treatment period. All mortalities occurred during the seven-day monitoring period, after completion of 7 days of repeated compound administration. Mortalities in the 8 µL M IP group began on day 8, with one mortality. Most mortalities occurred between days 9 and 10. All 8 µL M IP animals died or were euthanized by day 11. The survival curve for the 8 µL M IP group was significantly (p = 0.0002) different compared to that of the 8 µL S IP group. No mortalities occurred in the 4 µL M IP, 8 µL M PO or 8 µL S IP groups.

**Figure 4:**
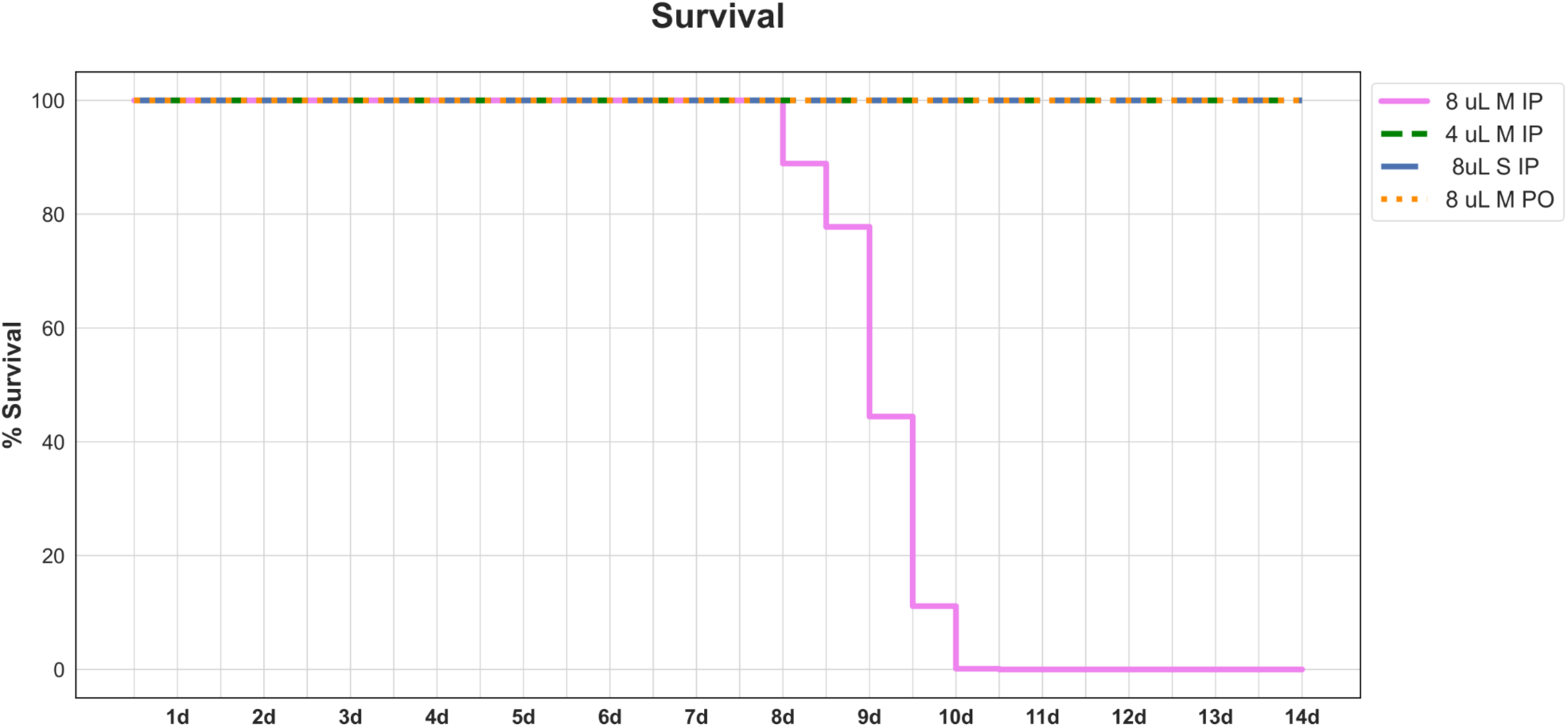
Survival Curve. Kaplan Meier plot represents percent survival throughout the 14-day experimental timeline. All animals in the 4 µL M IP, 8 µL S IP and 8 µL M PO groups survived to endpoint. Mortality in the 8 µL M IP group began on day 8.5, with the highest number of mortalities between day 9 and 10. Survival curve for the 8 µL M IP group, but no other group, was significantly (p = 0.0002) different than that of the 8 µL S IP group. All animals died by day 11. 8 µL M IP, day 1 *n* = 9, day 11 *n* = 0; 4 µL M IP, *n* = 10; 8 µL S IP, *n* = 10; 8 µL M PO, *n* = 7.

### Gross necropsy

At the monitoring period conclusion, or upon meeting euthanasia criteria, surviving animals were euthanized for gross necropsy (**Table 2 and Figure 5**). The 8µL M IP animals showed the most severe abnormalities on gross necropsy, including approximately 0.5 mL of serosanguinous fluid in the thoracic cavity (4/5 animals; **Figure 5A**); adhesions between the liver and diaphragm and between the stomach and liver (5/5 animals; adhesions also involved the kidney and pancreas/spleen in two mice; **Figure 5B-C**); a dark red to purple mottled appearance of the lungs (2/5 animals; in one additional animal the lungs were dark pink and sunk under the level of formalin); cystic masses containing oily fluid in the peri-reproductive fat (4/5 animals); and red-tan discoloration of the liver with rounded margins (4/5 animals; in an additional animal the surface of the liver was covered with stringy white material) (**Figure 5**). These findings were less prominent in the 4 µL M IP group, where only one mouse had adhesions in the abdomen and no mice had discoloration of the liver or serosanguinous fluid in the thoracic cavity. Mice in the 4 µL M IP group did have an abnormal dark red mottled appearance to the lungs in two cases and cystic masses in the peri-reproductive fat in all cases (5/5 animals). Mice in the 8 µL M PO and 8 µL S IP groups had no significant abnormalities on gross necropsy (5/5 animals).

**Figure 5:**
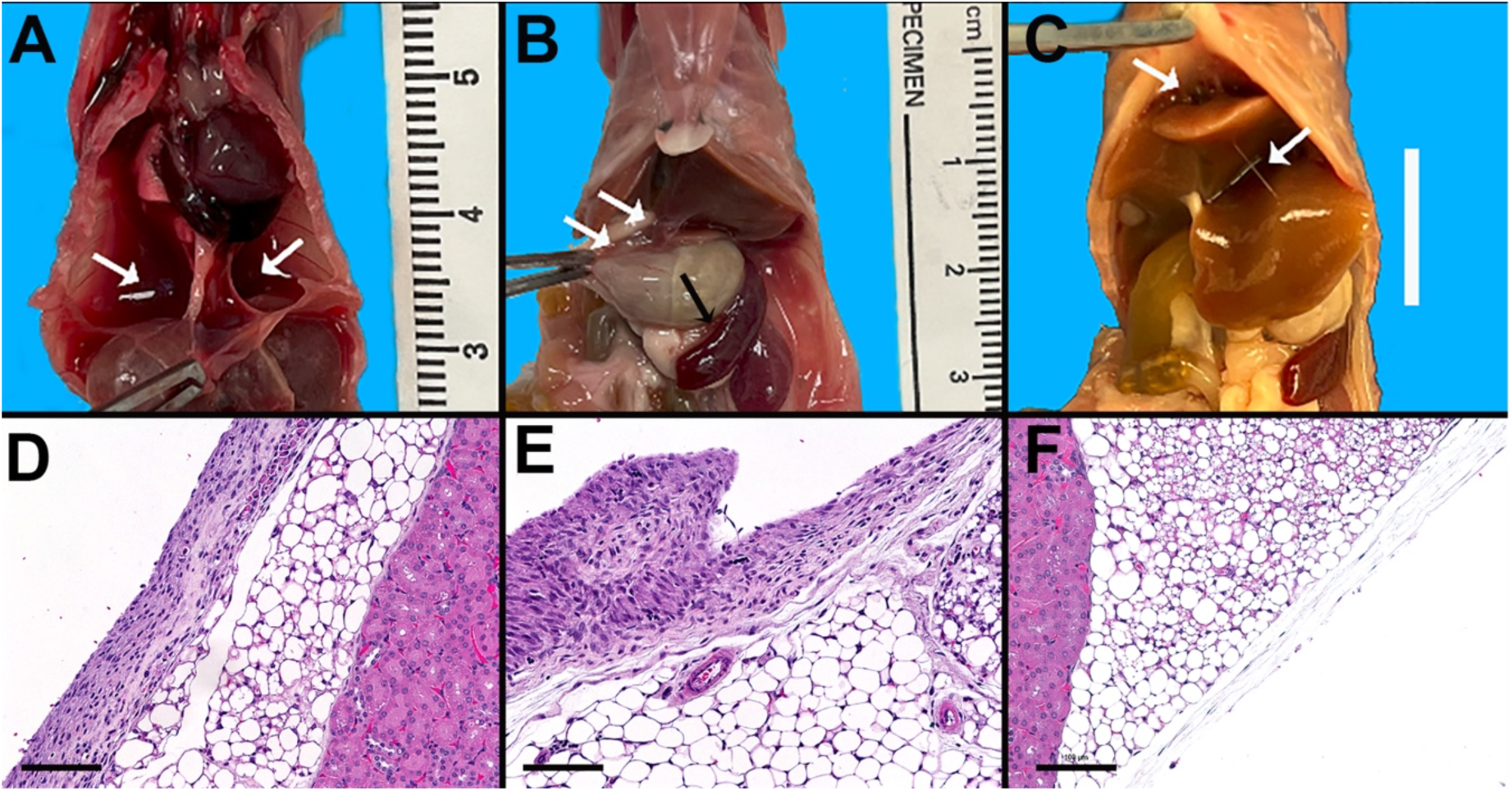
Gross Necropsy. A-C. Gross images of from 8 µL M IP mice, showing fluid within the thoracic cavity (A, white arrows); adhesions between the stomach and liver (B, white arrows) and between the stomach and enlarged spleen (black arrows) and adhesions between the liver and diaphragm and liver lobes (C, white arrows). D-F are histology images from the retroperitoneal tissues near the kidney from a (D) 8 µL M IP mouse; (E) 4 µL M IP mouse; and (F) 8 µL S IP mouse. Bar is 100 microns; hematoxylin and eosin (H&E).

**Table 2:** Gross Necropsy.

| Group | Gross lesions |  |  | Histologic lesions |  |  |  |  |
| --- | --- | --- | --- | --- | --- | --- | --- | --- |
|  | Serosanguinous<br>fluid thoracic<br>cavity | Abdominal<br>adhesions | Cystic masses<br>peri-reproductive<br>fat | Lung<br>hemorrhage<br>(0-4) | Lung<br>inflammation<br>(0-4) | Mediastinal<br>inflammation<br>(0-2) | Peritoneal<br>inflammation<br>(0-2) | Clear vacuoles<br>in lung<br>(Y/N) |
| 8 uL / g<br>Miglyol<br>IP | 4/5 | 5/5 | 5/5 | 0 | 2 | 1 | 2 | Y |
|  |  |  |  | 4 | 3 | 2 | 2 | Y |
|  |  |  |  | 3 | 2 | 2 | 2 | Y |
|  |  |  |  | 2 | 1 | 2 | 2 | Y |
|  |  |  |  | 1 | 2 | 2 | 2 | Y |
| 4 uL / g<br>Miglyol<br>IP | 0/5 | 1/5 | 5/5 | 2 | 1 | 0 | 1 | N |
|  |  |  |  | 1 | 2 | 0 | 1 | +/- |
|  |  |  |  | 1 | 1 | 0 | 1 | Y |
|  |  |  |  | 1 | 2 | 0 | 1 | N |
|  |  |  |  | 2 | 3 | 1 | 1 | Y |
| 8 uL / g<br>saline<br>IP | 0/2 | 0/2 | 0/2 | 1 | 0 | 0 | 0 | N |
|  |  |  |  | 1 | 1 | 0 | 0 | N |
| 8 uL / g<br>Miglyol<br>PO | 0/5 | 0/5 | 0/5 | 0 | 1 | 0 | 0 | N |
|  |  |  |  | 0 | 1 | 0 | 0 | N |
|  |  |  |  | 0 | 1 | 0 | 0 | N |
|  |  |  |  | 0 | 0 | 0 | 1 | N |
|  |  |  |  | 1 | 1 | 1* | 0 | N |

\*This mouse had minimal inflammation associated with arteritis in the mediastinal tissues, presumed to be an incidental finding.

### Histology

Terminal histopathology was also assessed in all animals (**Table 2 and Figure 6**). Variably severe acute pulmonary hemorrhage and inflammation was present in mice from the 8 µL M IP and the 4 µL M IP groups, with a statistically significant difference between all groups on Kruskal-Wallis test (p = 0.002 and p = 0.02, respectively; **Figure 6**). Acute hemorrhage was most severe (score of 4) in one mouse from the 8 µL M IP group. Inflammation was characterized by histiocytes with vacuolated cytoplasm and fewer neutrophils. Occasionally, inflammatory cells surrounded clear small circular structures (presumptive impression of a lipid droplet; **Figure 6**). Similar inflammation was observed throughout the mediastinal and mesenteric fat and along the serosal surfaces of the liver, kidney, and spleen and affecting the peritoneal fat in 8 µL M IP mice and to a much lesser extent the 4 µL M IP mice. The difference between all groups on Kruskal-Wallis test for both mediastinal and peritoneal inflammation was statistically significant (p < 0.0001; **Figure 6**). There was increased extramedullary hematopoiesis in the spleen in 3/5 8 µL M IP mice. The 8 µL M PO and 8 µL S IP groups presented with only minimal acute pulmonary hemorrhage (consistent with method of CO_2_ euthanasia), and no notable inflammation in the mediastinum and peritoneum.

**Figure 6:**
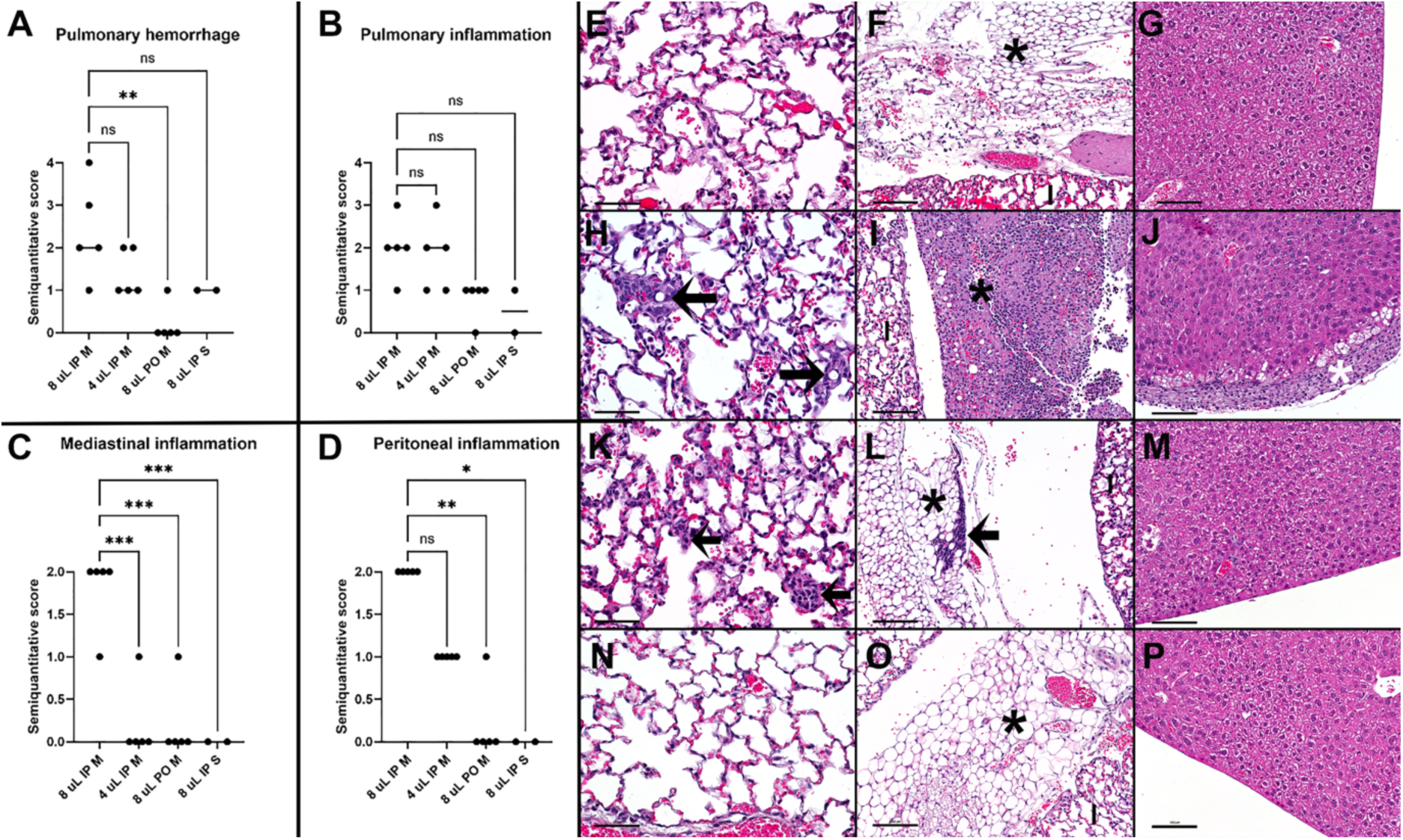
Histologic Scoring. Graphs of score distributions by group for pulmonary hemorrhage (A), pulmonary inflammation (B), mediastinal inflammation (C), and peritoneal inflammation (D). *** indicates p ≤ 0.001; ** indicates p ≤ 0.005; * indicates p = 0.01. E-P. Examples of histologic lesions observed in 8 µL S IP mice (top row, E-G); 8 µL M IP (second row, H-J), 4 µL M IP (third row, K-M), and 8 µL M PO (bottom row, N-P) for lung inflammation (left column, E, H, K, N); mediastinal inflammation (middle column, F, I, L, O); and peritoneal inflammation (right column, G, J, M, P). Black arrows in (H) indicate clear vacuoles surrounded by inflammation in the lung. In (K), black arrows indicate small foci of inflammatory cell infiltration without clear vacuoles. Black asterisks in F, I, L, and O indicates mediastinal fat, which in the 8 µL/g mouse (I) is effaced by inflammation, acute hemorrhage and fibrin. “l” indicates lung tissue. Black arrow in (L) indicates a mild accumulation of lymphocytes, which can be an incidental finding in mice. In (J), the white asterisk indicates inflammation adjacent to the liver margin in an 8 µL M IP mouse, which is not observed in the 4 µL M IP and 8 µL S IP mice. Bar in E H, K, and N is 50 microns; all other images bar is 100 microns. H&E.

### Complete Blood Count

CBC results assessed general animal well-being and disease status resulting from repeated pharmaceutical-grade MCTs or saline administration (**Table 3**). 8 µL M IP animals showed markedly elevated absolute WBC counts characterized by a neutrophilia and monocytosis as compared to all other groups in our study, as well as the mean ± standard deviation for C57BL/6J males (The Jackson Laboratory, 2020). The 8 µL M IP group also showed slightly elevated MCH and MCHC, and slightly depleted RBC, HCT and reticulocyte count, as compared to data from Jackson Laboratory and other treatment groups. Of all cell types analyzed, only RBC count in the 8 µL M IP group was significantly different compared to the 8 µL S IP group (p = 0.02). However, this may be attributed to the low sample size (n = 2) of blood collected from the 8 µL S IP group. The RBC and HCT in 8 µL M IP group were also significantly different (p = 0.04 and 0.03, respectively) than the 4 µL M IP group but not the 8 µL M PO group. There was no significant difference in any other CBC metric.

**Table 3:**
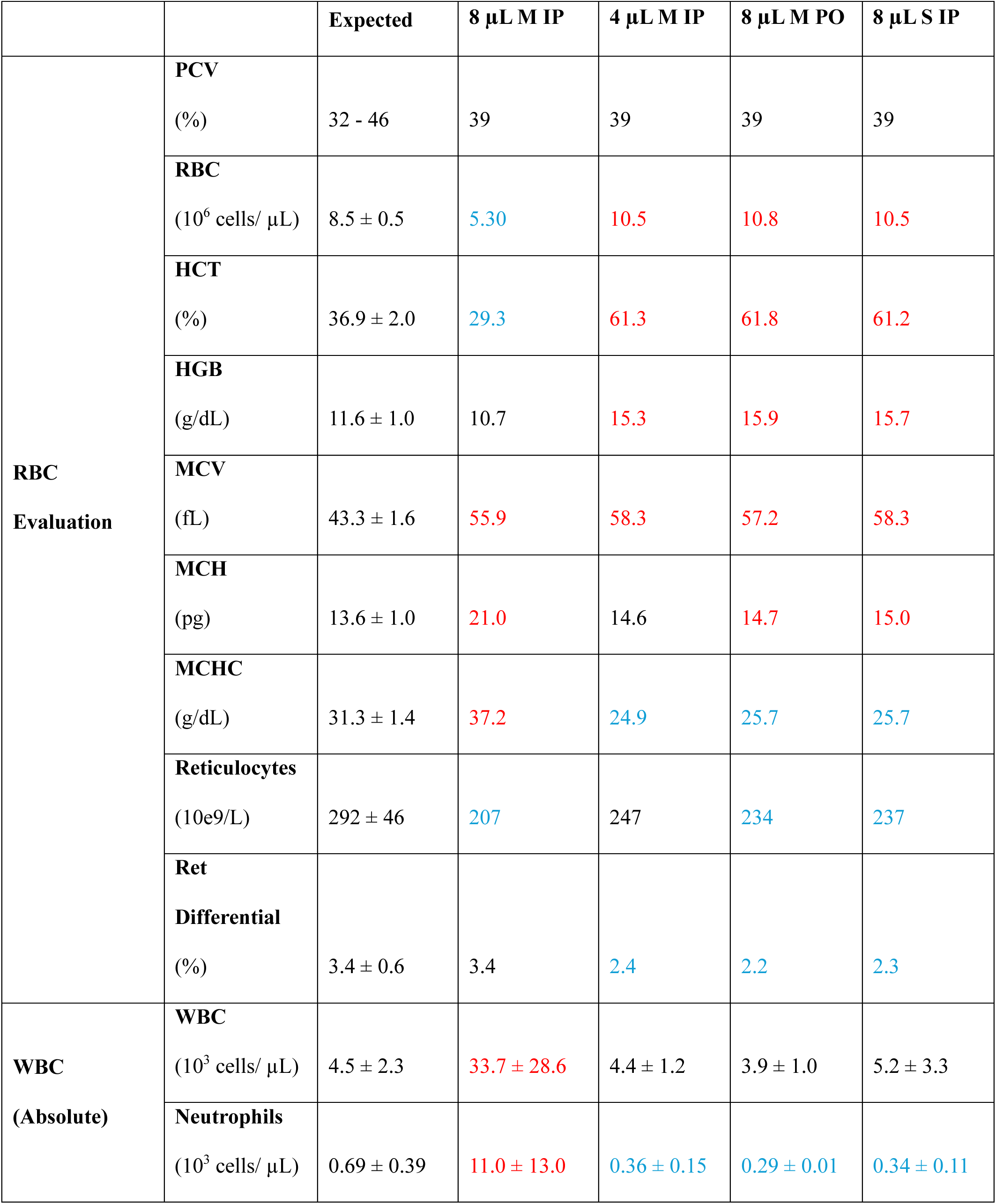
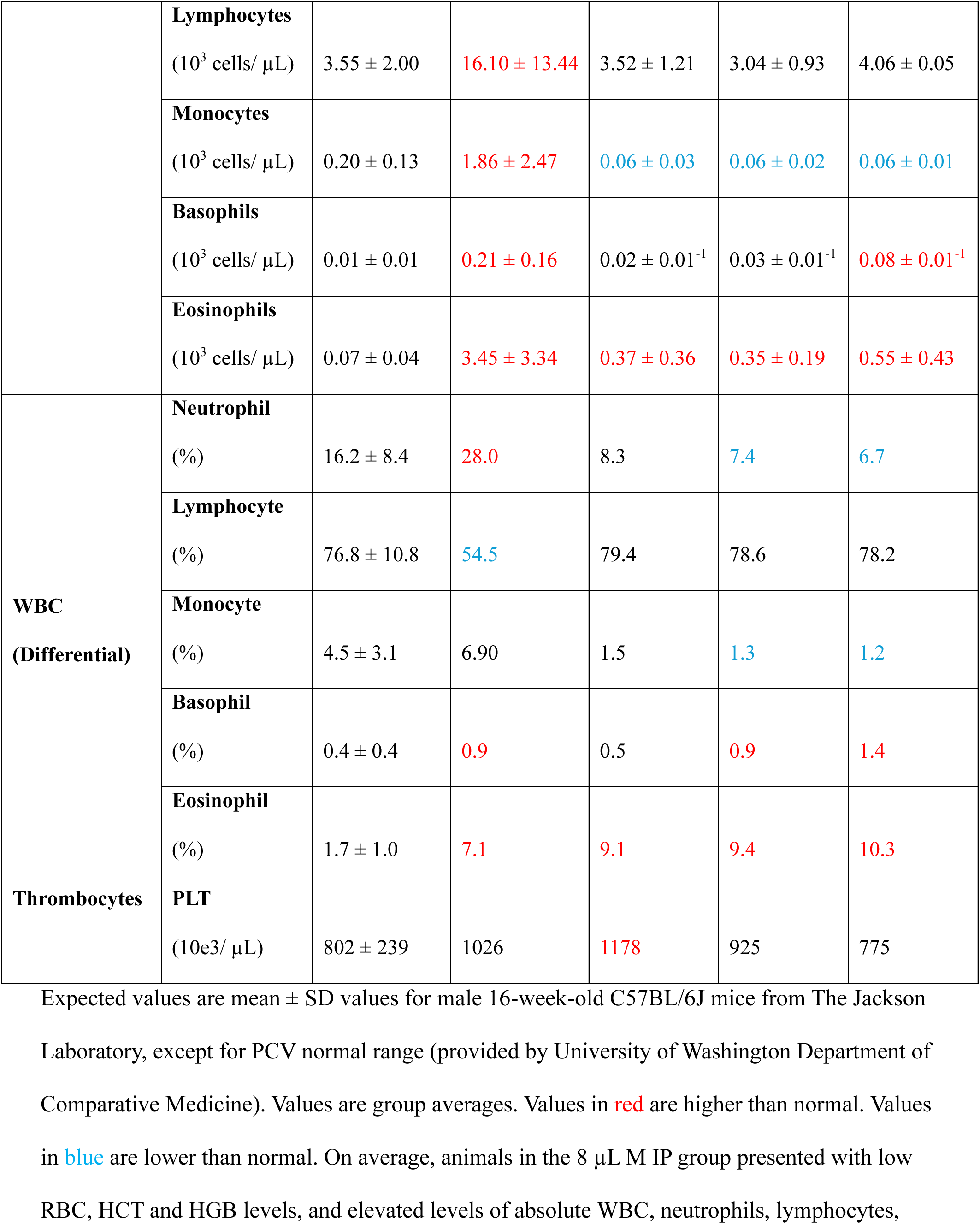
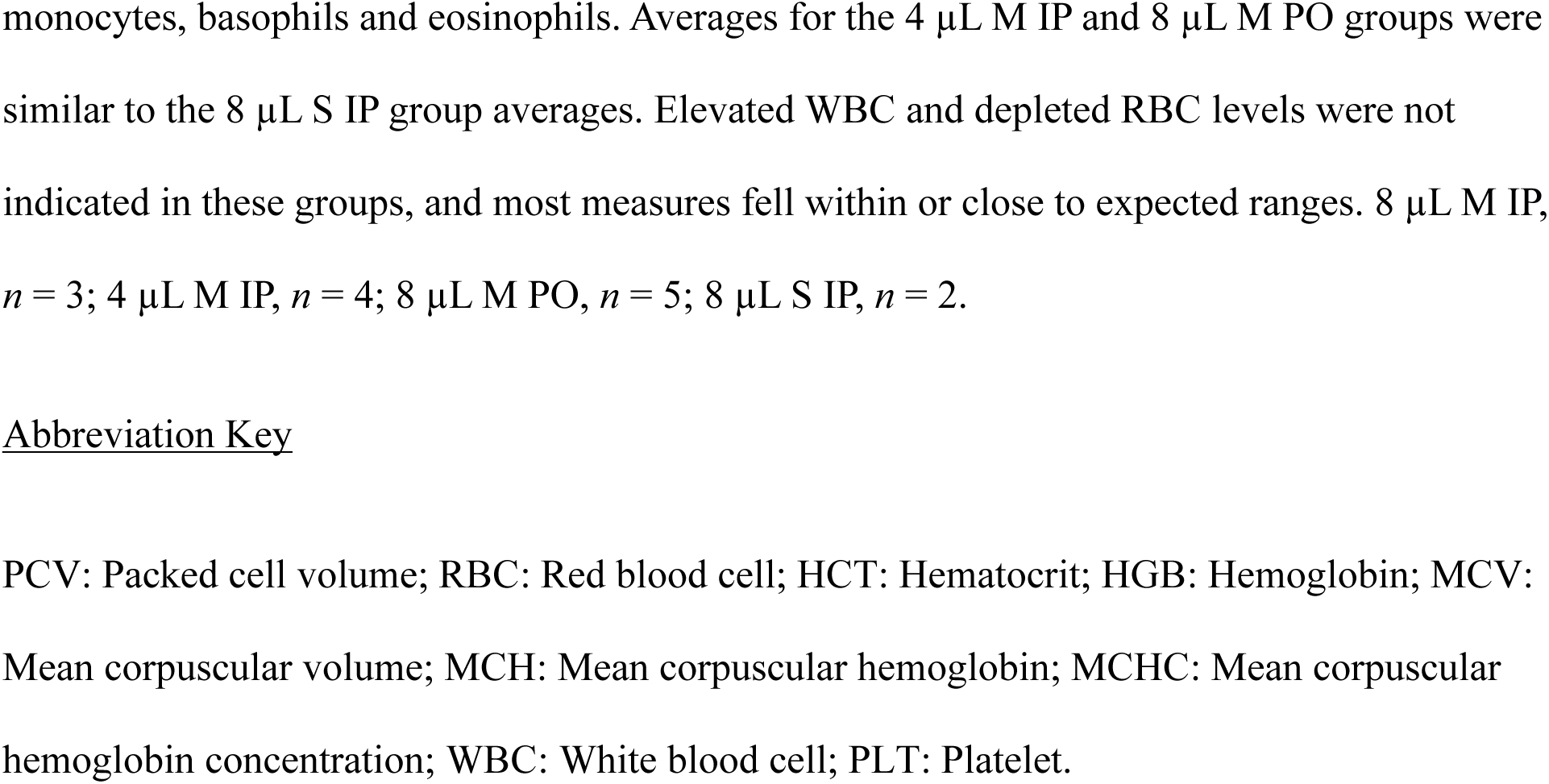
Complete Blood Count.

| | | Expected | 8 μL M IP | 4 μL M IP | 8 μL M PO | 8 μL S IP |
| --- | --- | --- | --- | --- | --- | --- |
| **RBC**<br>**Evaluation** | **PCV**<br>(%) | 32 - 46 | 39 | 39 | 39 | 39 |
| | **RBC**<br>($10^6$ cells/ μL) | $8.5 \pm 0.5$ | 5.30 | 10.5 | 10.8 | 10.5 |
| | **HCT**<br>(%) | $36.9 \pm 2.0$ | 29.3 | 61.3 | 61.8 | 61.2 |
| | **HGB**<br>(g/dL) | $11.6 \pm 1.0$ | 10.7 | 15.3 | 15.9 | 15.7 |
| | **MCV**<br>(fL) | $43.3 \pm 1.6$ | 55.9 | 58.3 | 57.2 | 58.3 |
| | **MCH**<br>(pg) | $13.6 \pm 1.0$ | 21.0 | 14.6 | 14.7 | 15.0 |
| | **MCHC**<br>(g/dL) | $31.3 \pm 1.4$ | 37.2 | 24.9 | 25.7 | 25.7 |
| | **Reticulocytes**<br>($10^9$/L) | $292 \pm 46$ | 207 | 247 | 234 | 237 |
| | **Ret**<br>**Differential**<br>(%) | $3.4 \pm 0.6$ | 3.4 | 2.4 | 2.2 | 2.3 |
| **WBC**<br>**(Absolute)** | **WBC**<br>($10^3$ cells/ μL) | $4.5 \pm 2.3$ | $33.7 \pm 28.6$ | $4.4 \pm 1.2$ | $3.9 \pm 1.0$ | $5.2 \pm 3.3$ |
| | **Neutrophils**<br>($10^3$ cells/ μL) | $0.69 \pm 0.39$ | $11.0 \pm 13.0$ | $0.36 \pm 0.15$ | $0.29 \pm 0.01$ | $0.34 \pm 0.11$ |
| | **Lymphocytes**<br>(10<sup>3</sup> cells/ μL) | 3.55 ± 2.00 | 16.10 ± 13.44 | 3.52 ± 1.21 | 3.04 ± 0.93 | 4.06 ± 0.05 |
| | **Monocytes**<br>(10<sup>3</sup> cells/ μL) | 0.20 ± 0.13 | 1.86 ± 2.47 | 0.06 ± 0.03 | 0.06 ± 0.02 | 0.06 ± 0.01 |
| | **Basophils**<br>(10<sup>3</sup> cells/ μL) | 0.01 ± 0.01 | 0.21 ± 0.16 | 0.02 ± 0.01<sup>-1</sup> | 0.03 ± 0.01<sup>-1</sup> | 0.08 ± 0.01<sup>-1</sup> |
| | **Eosinophils**<br>(10<sup>3</sup> cells/ μL) | 0.07 ± 0.04 | 3.45 ± 3.34 | 0.37 ± 0.36 | 0.35 ± 0.19 | 0.55 ± 0.43 |
| **WBC**<br>**(Differential)** | **Neutrophil**<br>(%) | 16.2 ± 8.4 | 28.0 | 8.3 | 7.4 | 6.7 |
| | **Lymphocyte**<br>(%) | 76.8 ± 10.8 | 54.5 | 79.4 | 78.6 | 78.2 |
| | **Monocyte**<br>(%) | 4.5 ± 3.1 | 6.90 | 1.5 | 1.3 | 1.2 |
| | **Basophil**<br>(%) | 0.4 ± 0.4 | 0.9 | 0.5 | 0.9 | 1.4 |
| | **Eosinophil**<br>(%) | 1.7 ± 1.0 | 7.1 | 9.1 | 9.4 | 10.3 |
| **Thrombocytes** | **PLT**<br>(10e3/ μL) | 802 ± 239 | 1026 | 1178 | 925 | 775 |

Expected values are mean ± SD values for male 16-week-old C57BL/6J mice from The Jackson Laboratory, except for PCV normal range (provided by University of Washington Department of Comparative Medicine). Values are group averages. Values in red are higher than normal. Values in blue are lower than normal. On average, animals in the 8 μL M IP group presented with low RBC, HCT and HGB levels, and elevated levels of absolute WBC, neutrophils, lymphocytes,
monocytes, basophils and eosinophils. Averages for the 4 μL M IP and 8 μL M PO groups were similar to the 8 μL S IP group averages. Elevated WBC and depleted RBC levels were not indicated in these groups, and most measures fell within or close to expected ranges. 8 μL M IP, $n = 3$ ; 4 μL M IP, $n = 4$ ; 8 μL M PO, $n = 5$ ; 8 μL S IP, $n = 2$ .

## DISCUSSION

This study assesses the safety of escalating concentrations of repeated IP versus PO administration of pharmaceutical-grade MCTs in a cohort of experimentally naïve C57BL/6J male mice to understand whether this vehicle could be associated with any untoward effects on animal well-being and/or survival. The acute oral LD_50_ for the pharmaceutical-grade MCTs presently employed is reported as > 5.0 g/kg in rats (Spectrum Chemical, 2022), but there is no published LD_50_ for IP administration in mice. Here, repeated IP administration of a high dose (8 µL/g BW or 7.56 g/kg) of 100% pharmaceutical-grade MCTs led to severe pathology and 100% mortality in our group of male mice. This adverse response was not seen in animals receiving repeated IP administration of a lower dose (4 µL/g BW or 3.78 g/kg) or the same high dose PO. As such, for repeated IP administration our study determines an acutely toxic and fatal dose (8 µL/g) and a non-fatal dose (4 µL/g). The IP LD_50_ for our pharmaceutical-grade MCTs likely resides between these two doses. Our study did not determine a no observed adverse effect level (NOAEL) for repeated IP pharmaceutical-grade MCTs, because animals receiving the 4 µL/g dose still presented with mild lesions and transient weight loss without clinical decline. In contrast, we found that repeated administrations of the 8 µL/g dose of pharmaceutical-grade MCTs via the oral route was tolerable, suggesting a NOAEL for PO pharmaceutical-grade MCTs.

Our histopathological findings suggest that repeated administration of high dose pharmaceutical-grade MCTs leads to abdominal accumulation of more lipid than the body can process, potentially causing systemic inflammation, pulmonary lesions, and pleural effusion. Cystic lesions and variably severe inflammation, without pleural effusion, in the 4 µL/g M IP group suggests a dose-dependent effect of pharmaceutical-grade MCTs toxicity, where repeated IP delivery of 4 µL/g BW is tolerated but 8 µL/g overwhelms the system. Elevated absolute WBCs seen in the 8 µL/g M IP but not 4 µL/g M IP mice are further evidence of a dose-dependent systemic inflammatory response. Lack of these effects with PO administration suggests a route-dependent effect. Absence of lesions in animals receiving saline IP suggests the IP route itself is not responsible for the findings in the 8 µL/g M IP and 4 µL/g M IP groups.

MCTs used in clinical practice is further evidence of a dose- and route-dependent effect for pharmaceutical-grade MCTs. MCTs are administered orally for nutritional support in gastrointestinal (GI) disorders. Containing only saturated fatty acids, they are rapidly absorbed passively from the GI tract into the bloodstream via the portal vein. In digestion, MCTs require minimal packaging, making them an effective source of rapid caloric intake. This makes MCTs an appealing treatment for malabsorption disorders, including pancreatic insufficiency, and following gastrectomy or small bowel resection (Shah & Limketkai, 2017). So, it is unsurprising that we did not see major adverse responses to PO administration here.

Our present study illustrates no adverse effect of repeated high dose pharmaceutical-grade MCTs orally administered in male mice, even at a dose well-above the rat LD_50_. When administered orally, MCTs are broken down into monocarboxylic saturated fatty acids (MCFAs), which are then absorbed through the portal vein into the bloodstream and lymphatic system (You, 2011). Metabolism of MCFAs by hepatic mitochondria produces high levels of acetyl co-enzyme A, stimulating ketone body production (Traul, 1999; Lemarié, 2015). A growing body of evidence supports the use of MCTs in ketogenic metabolic therapy for treatment of neurological disorders including epilepsy (Augstin, 2018; Wláz, 2014), neurodegeneration (Joniec-Maciejak, 2018; Zhao, 2012), and others (Altinoz, 2020; Augur, 2018; Hecker, 2014; Seyfried, 2019; Winter, 2017). Our present study confirms that repeated oral administration of MCT-containing compounds is well-tolerated in mice. Interestingly, oral absorption of long chain-versus MCT-conjugated antiseizure medicines can dramatically change drug bioavailability, with food intake influencing drug effects (Dahan, 2008). Further investigation into the benefit of orally administered MCT-formulated therapeutics in neurological disease models is needed.

There is little literature regarding the processing of MCTs in the peritoneum. IP administration of MCTs bypasses GI mechanisms entirely. This includes gastric acid lipase, which hydrolyzes triglycerides into diglycerides and fatty acids, and pancreatic fluid and bile that initiate further hydrolysis and stabilization of smaller emulsions (Yáñez, 2011). IP-delivered MCTs likely cannot be broken down into MCFAs. Large MCT molecules may accumulate into cystic lesions, initiating inflammation. If taken up via diffusion into highly permeable lymphatic capillaries, MCTs could cause blockages or be deposited elsewhere in the body, including the thoracic cavity as we hypothesize occurred here. Further evidence for this hypothesis is reported adverse events of repeated IP dosing with methylcellulose (MC), another commonly-used hydrophobic excipient in pharmaceutical sciences research. Mirroring our findings, Meeker et al. found lesions in the peritoneal cavity and lungs, and elevated WBC count in animals receiving repeated MC dosing via the IP, but not PO, route (Meeker, 2019). Thus, repeated IP administration of high concentrations of hydrophobic excipients in mice likely induces significant thoracic accumulation that may negatively affect physiological outcomes, potentially confounding investigational pharmaceutical compound studies.

The clinical presentations and gross and histologic lesions seen here suggest that repeated administration of pharmaceutical-grade MCTs by the IP route may cause systemic toxicity. However, this effect is dose-dependent, as a lower dose delivered IP, as well as a high dose delivered orally, were generally well-tolerated. Our reported results clearly demonstrate that the best use of pharmaceutical-grade MCTs and similar excipients is restricted to low doses or oral administration. This study may complicate the use of MCTs as a formulation vehicle because compound absorption into the bloodstream is much more efficient via the IP as compared to PO route (Al Shoyaib, 2020). Nemes et al. found bioavailability of some compounds to be as much as 6-fold higher and 4-fold faster via IP administration (Nemes, 2000). Long chain triglyceride (LCT)-conjugated compounds may be more orally bioavailable. Dahan et al. found that a phospholipid-valproic acid conjugate (DP-VPA) had three-fold higher bioavailability in LCT as compared to MCT solution (Dahan, 2008). However, PO administration may still require either a larger dose of excipient or a higher concentration of drug to achieve necessary blood levels of drug. A caveat of increasing excipient dose is previous adverse reactions in rodents receiving PO administration of >10 µL of Miglyol/g BW (Thackaberry, 2013). On the other hand, some investigational compounds may precipitate out of solution at the higher concentrations needed to achieve effective plasma and target organ concentrations *in vivo*. This is particularly challenging in CNS disease studies, where drug delivery across the blood-brain barrier is further limited. Due to their rapid absorption and CNS bioavailability, MCTs make a good vehicle in these studies. In this case, if IP administration is necessary, drug concentration should be maximized to reduce vehicle dosage and associated adverse effects.

The findings presented here provide important insight regarding the safest usage of a widely used formulation vehicle for preclinical drug discovery research. It is important to note certain limitations of this present study. Our study shows dose- and route-dependent effects of repeated administration. In support of this, previous work using lower and/or single doses of MCT excipients via the IP route, and administration via the sub-cutaneous, intravenous and oral route in various species have proven safe (Baazm, 2021; Buss, 2018; Bunaciu, 2011; Rochette, 2008). Conversely, longer or more intensive dosing via the PO route or even lower doses via the IP route may cause more severe pathology than seen here. Histologic lesions observed in the 4 µL M IP animals, despite lack of clinical decline may indicate there is a cumulative threshold dose above which MCTs toxicity occurs. Longer and/or more intensive dosing may reach this cumulative threshold at lower doses. It is also important to note that our study used only male mice, and female animals may respond differently.

In summary, our findings suggest that although many uses of pharmaceutical-grade MCTs are safe in rodents, repeated administration of high dosages via the IP route may lead to pathology and/or mortality. Importantly, histologic lesions may be present even without overt physical or behavioral changes. When using pharmaceutical-grade MCTs as a vehicle, researchers should consider which method of administration is most effective, while also minimizing risk of adverse events. Things to consider in this evaluation are the MCT dose needed to deliver the desired drug, its relative absorption via the IP vs PO route, duration and frequency of the administration paradigm, and the potential impact that animals’ stress may have on experimental results. Finally, this endeavor emphasizes the importance of including control and sham groups in pharmacological and disease model studies to account for unanticipated effects of vehicle solutions. Additional study of MCT-containing formulation vehicles in mice may further elucidate causes of clinical decline and pathology and help optimize dosing regimens.

## ACKNOWLEDGMENTS

- Dr. Thea L. Brabb
- Wenxuan Xiong
- University of Washington Department of Comparative Medicine Veterinary Diagnostic Laboratory and Histological and Imaging Core

Funding for this project was provided by NIH/NINDS grant NS123195 (JR Weinstein)

## DATA AVAILABILITY STATEMENT

The authors declare that all the data supporting the findings of this study are available within the paper and its Supplemental Data.

## AUTHORSHIP CONTRIBUTIONS

*Participated in research design*: Adler-Wachter, Barker-Haliski, and Weinstein.

*Conducted experiments*: Adler-Wachter, Tsai, Schweitzer, and Snyder.

*Contributed new reagents or analytical tools*: Snyder, and Weinstein.

*Performed data analysis*: Adler-Wachter, and Snyder.

*Wrote or contributed to the writing of the manuscript*: Adler-Wachter, Tsai, McDonough, Snyder, Barker-Haliski, and Weinstein.

---

**Supplemental Table 1.**

|  | Lung<br>hemorrhage | Lung<br>inflammation | Mediastinal<br>inflammation | Peritoneal<br>inflammation |
| --- | --- | --- | --- | --- |
| 0 | None | None (minimal<br>lymphocytic<br>aggregates may<br>be present) | None (minimal<br>lymphocytic<br>aggregates may<br>be present) | None (minimal<br>lymphocytic<br>aggregates may<br>be present) |
| 1 | Minimal<br>multifocal<br>alveolar<br>hemorrhage,<br>affecting <5% of<br>lung | Minimal, few<br>foci of<br>inflammation, 5<br>or fewer | Minimal to mild | Minimal to mild |
| 2 | Mild multifocal,<br>5-19% of lung | Mild multifocal<br>foci of<br>inflammation | Moderate to<br>severe, may<br>involve serosal<br>surface of heart | Moderate to<br>severe, involves<br>serosal surface<br>of liver, kidney,<br>and mesenteric<br>fat |
| 3 | Moderate<br>multifocal to<br>coalescing, 20-50% | Moderate,<br>multifocal to<br>coalescing | n/a | n/a |
| 4 | Severe, >50% | Severe,<br>multifocal to<br>coalescing | n/a | n/a |

## FOOTNOTES

* Funding for this project was provided by NIH/NINDS grant NS123195 (JRW).

## REFERENCES

Al Shoyaib A, Archie SR and Karamyan VT (2020) Intraperitoneal route of drug administration: should it be used in experimental animal studies? Pharm Res 37:12. DOI: 10.1007/s11095-019-2745-x.

Altinoz MA, Ozpinar A and Seyfried TN (2020) Caprylic (octanoic) acid as a potential fatty acid chemotherapeutic for glioblastoma. Prostaglandins Leukot Essent Fatty Acids 159:102142. DOI: 10.1016/j.plefa.2020.102142.

Augur ZM, Doyle CM, Li M, Mukherjee P and Seyfried TN (2018) Nontoxic targeting of energy metabolism in preclinical VM-M3 experimental glioblastoma. Front Nutr 5:19. DOI: 10.3389/fnut.2018.00091.

Augustin K, Khabbush A, Williams S, Eaton S, Orford M, Cross JH, Heals SJR, Walker MC and Williams RSB (2018) Mechanisms of action for the medium-chain triglyceride ketogenic diet in neurological metabolic disorders. Lancet Neurol 17:84–93.

Baazm M, Behrens V, Beyer C, Nikoubashman O and Zendedel A (2021) Regulation of inflammasomes by application of omega-3 polyunsaturated fatty acids in a spinal cord injury model. Cells 10. DOI: 10.3390/cells10113147.

Bunaciu RP, Tharappel JC, Lehmler HJ, Lee EY, Robertson LW, Bruckner GC, Spear BT and Glauert HP (2011) Role of oil vehicle on hepatic cell proliferation in PCB-treated rats. J Environ Pathol Toxicol Oncol 30:273–282.

Burkholder T, Foltz C, Karlsson E, Linton GC and Smith JM (2012) Health evaluation of experimental laboratory mice. Curr Protoc Mouse Biol 2:145–165. DOI: 10.1002/9780470942390.mo110217.

Buss N, Ryan P, Baughman T, Roy D, Patterson C, Gordon C and Dixit R (2018) Nonclinical safety and pharmacokinetics of Miglyol 812: A medium chain triglyceride in exenatide once weekly suspension. J App Toxicol 38:1293–1301. DOI: 10.1002/jat.3640.

Dahan A, Duvdevani R, Shapiro I, Elmann A, Finklestein E and Hoffman A (2008) Absorption of phospholipid prodrugs: in vivo and in vitro mechanistic investigation of trafficking of a lecithin- valproic acid conjugate following oral administration. J Control Release 126:1–9. DOI: 10.1016/j.jconrel.2007.10.025.

Healing G, Sulemann T, Cotton P, Harris J, Hargreaves A, Finney R, Kirk S, Schramm C, Garner C, Pivette P and Burdett L (2016) Safety data on 19 vehicles for use in 1 month oral rodent pre- clinical studies: administration of hydroxypropyl-ß-cyclodextrin causes toxicity. J Appl Toxicol 36:140–150. DOI: 10.1002/jat.3155.

Hecker M, Sommer N, Voigtmann H, Pak O, Mohr A, Wolf M, Vadász I, Herold S, Weissmann N, Morty RE, Seegeer W and Mayer K (2014) Impact of short- and medium-chain fatty acids on mitochondrial function in severe inflammation. JPEN J Parenter Enteral Nutr 38:587–594. DOI: 10.1177/0148607113489833.

Joniec-Maciejak I, Wawer A, Turzyńska D, Sobolewska A, Maciejak P, Szyndler J, Mirowska-Guzel D and Płaźnik A (2018) Octanoic acid prevents reduction of striatal dopamine in the MPTP mouse model of Parkinson’s disease. Pharmacol Rep 70:988–992. DOI: 10.1016/j.pharep.2018.04.008.

Lee RD, Chen YJ, Nguyen HM, Singh L, Dietrich CJ, Pyles BR, Cui Y, Weinstein JR and Wulff H (2024) Repurposing the K_Ca_3.1 blocker senicapoc for ischemic stroke. Transl Stroke Res 15:518–532. DOI 10.1007/s12975-023-01152-6.

Lemarié F, Beauchamp E, Legrand P and Rioux V (2016) Revisiting the metabolic and physiological functions of caprylic acid (C8:0) with special focus on ghrelin octanoylation. Biochimie 120:40–48. DOI: 10.1016/j.biochi.2015.08.002.

Meeker S, Beckman M, Knox KM, Treuting PM and Barker-Haliski M (2019) Repeated intraperitoneal administration of low-concentration methylcellulose leads to systemic histologic lesions without loss of preclinical phenotype. J Pharmacol Exp Ther 371:25–35. DOI: 10.1124/jpet.119.257261.

National Academy of Science. 2011. Guide for the Care and Use of Laboratory Animals 8th ed. The National Academic Press.

Nemes KB, Abermann M, Bojti E, Grézal G, Al-Behaisi S and Klembovich I (2000) Oral, intraperitoneal and intravenous pharmacokinetics of deramciclane and its *N*-desmethyl metabolite in the rat. J Pharm Pharmacol 52:47–51.

Robaina Cabrera CL, Keir-Rudman S, Horniman N, Clarkson N and Page C (2021) The anti-inflammatory effects of cannabidiol and cannabigerol alone, and in combination. Pulm Pharmacol Ther 69:102047. DOI: 10.1016/j.pupt.2021.102047.

Rochette A, Hocquet AF, Dadure C, Boufroukh D, Raux O, Lubrano JF, Bringuier S and Capdevila X (2008) Avoiding propofol injection pain in children: a prospective, randomized, double-blinded, placebo-controlled study. Br J Anesth 101:390–394. DOI: 10.1093/bja/aen169.

Sellers RS, Antman M, Phillips J, Khan KN and Furst SM (2008) Effects of Miglyol 812 on rats after 4 weeks of gavage as compared with methylcellulose/tween 80. Drug Chem. Toxicol 28:423–432. DOI: 10.1080/01480540500262839.

Seyfried TN, Shelton L, Arismendi-Morillo G, Kalamian M, Elsakka A, Maroon J and Mukherjee P (2019) Provocative question: should ketogenic metabolic therapy become the standard of care for glioblastoma? Neurochem Res 44:2392–2404. DOI: 10.1007/s11064-019-02795-4.

Shah ND, and Limketkai BN (2017) The use of medium-chain triglycerides in gastrointestinal disorders. Pract Gastroenterol 120:20–28.

Singh Negi, J. 2019. Chapter 6 – Nanolipid materials for drug delivery systems: A comprehensive review. In: Mohapatra SS, Ranjan S, Dasgupta N, Kumar Mishra R, Thomas S, editors. Characterization and Biology of Nanomaterials for Drug Delivery. Elsevier (S&T), p. 137–163. DOI: 10.1016/C2017-0-00272-0.

Thackaberry EA (2013) Vehicle selection for nonclinical oral safety studies. Expert Opin Drug Metab Toxicol 9:1635–1646. DOI: 10.1517/17425255.2013.840291.

Traul KA, Driedger A, Ingle DL and Nakhasi D (1999) Review of the toxicologic properties of medium-chain triglycerides. Food Chem Toxicol 38:79–98.

Winter SF, Loebel F and Dietrich J (2017) Role of ketogenic metabolic therapy in malignant glioma: a systematic review. Crit Rev Oncol Hematol 112:41–58. DOI: 10.1016/j.critrevonc.2017.02.016.

Wlaź P, Socała K, Nieoczym D, Żarnowski T, Żarnowska I, Czuczwar SJ and Gasior M (2015) Acute anticonvulsant effects of capric acid in seizure tests in mice. Prog Neuropsychopharmacol Biol Psychiatry 57:110–116. DOI: 10.1016/j.pnpbp.2014.10.1013.

Yáñez JA, Wang SW, Knemeyer IW, Wirth MA and Alton KB (2011) Intestinal lymphatic transport for drug delivery. Adv Drug Deliv Rev 63:923–942. DOI: 10.1016/j.addr.2011.05.019.

You YQ, Ling PR, Qu JZ and Bistrian BR (2011) Effects of medium-chain triglycerides, long-chain triglycerides, or 2-monododecanoin on fatty acid composition in the portal vein, intestinal lymph, and systemic circulation in rats. JPEN J Parenter Enteral Nutr 32:169–175. DOI: 10.1177/0148607108314758.

Zhao W, Varghese M, Vempati P, Dzhun A, Cheng A, Wang J, Lange D, Bilski A, Faravelli I and Pasinetti GM (2012) Caprylic triglyceride as a novel therapeutic approach to effectively improve the performance and attenuate the symptoms due to the motor neuron loss in ALS disease. PLoS One 7:e49191.

